# GBM model refinement with literature curation, rule-based NLP, and LLMs

**DOI:** 10.1101/2025.03.27.645730

**Authors:** Niloofar Arazkhani, Haomiao Luo, Difei Tang, Brent Cochran, Natasa Miskov-Zivanov

## Abstract

In this work, our goal was twofold: (1) improve an existing glioblastoma multiforme (GBM) executable mechanistic model and (2) evaluate the effectiveness traditional natural language processing (NLP) pipeline and the generative AI approach in the process of model improvement. We used a suite of graph metrics and tools for interaction filtering and classification to collect data and conduct the analysis. Our results suggest that a more comprehensive literature search is necessary to collect enough information through automated paper retrieval and interaction extraction. Additionally, we found that graph metrics present a promising approach for model refinement, as they can provide useful insights and guidance when selecting new information to be added to a mechanistic model.

## I. Introduction

Different glioblastoma (GBM) stem cell lines can exhibit distinct genetic and molecular profiles, influencing their response to treatments. We have previously built an executable, mechanistic model of GBM stem cells to study the dynamic response of three different cell lines to a range of kinase inhibitors [1, 2]. This GBM model was developed by combining experimental data, information from databases and literature, and expert knowledge. It incorporates key GBM pathways and their cross talk, as well as intertwined feedforward and feedback loops, from receptor activation, through intermediate signaling molecules, to downstream transcriptional events that govern cell fate. The model has 410 elements, with 12 receptors, 129 proteins, 4 chemicals, 130 genes, and 9 biological processes. Receptors that are included in the model, through which input signals get propagated to cell, are EGFR, Insulin receptor PDGFRA, TGFBR, TNFR, VEGFR, and the biological processes modeled are apoptosis, cell cycle progression, DNA damage, hypoxia, neuronal differentiation, proliferation, protein synthesis, and stemness. Element interactions and regulatory functions are inferred from the information in literature and guided by expert knowledge, while mutations and element state assignments match experimental data. The model was simulated to obtain both transient and steady state values for each of the three cell lines before and after the treatment. These scenarios test the effectiveness of individual kinase inhibitors, a class of drugs that target specific kinases involved in cancer cell survival, as well as their combinations. The model makes predictions about temporal changes in proteins and genes, and cellular processes under these different scenarios.

Model was verified using literature and databases (structure), and by comparing simulation results with experimental data from wet lab studies (dynamic). While the model reproduces many of the experimental results, it still does not match all of them. Therefore, we explored whether the model could be improved with a fully automated approach. Such a workflow changes two aspects of the model, its structure, which is a directed network of connected nodes, and its state transition function which is comprised of individual node update rules. In our previous work we have also demonstrated that literature search queries and automated algorithms influence on identifying most accurate executable models [3, 4].

Here, we investigated: (1) the role of literature selected by an expert when refining the model; (2) the utility of traditional rule-based natural language processing (NLP) and new large language models (LLMs) in an automated flow from selected literature to model enhancement. Specifically, we focus on differences in node and network features between the manually built model and the NLP and LLM outputs.

## II. Methods

As illustrated in Figure 1, our workflow starts with an existing mechanistic model and a list of papers that an expert has been collecting in SciWheel [5], a cloud-based reference management tool over the course of a multi-year long project. We processed the collected papers using two approaches independently: a traditional NLP approach and a generative AI approach. We chose to use two complementary approaches due to the limitations of each when used alone.

**Figure 1.**
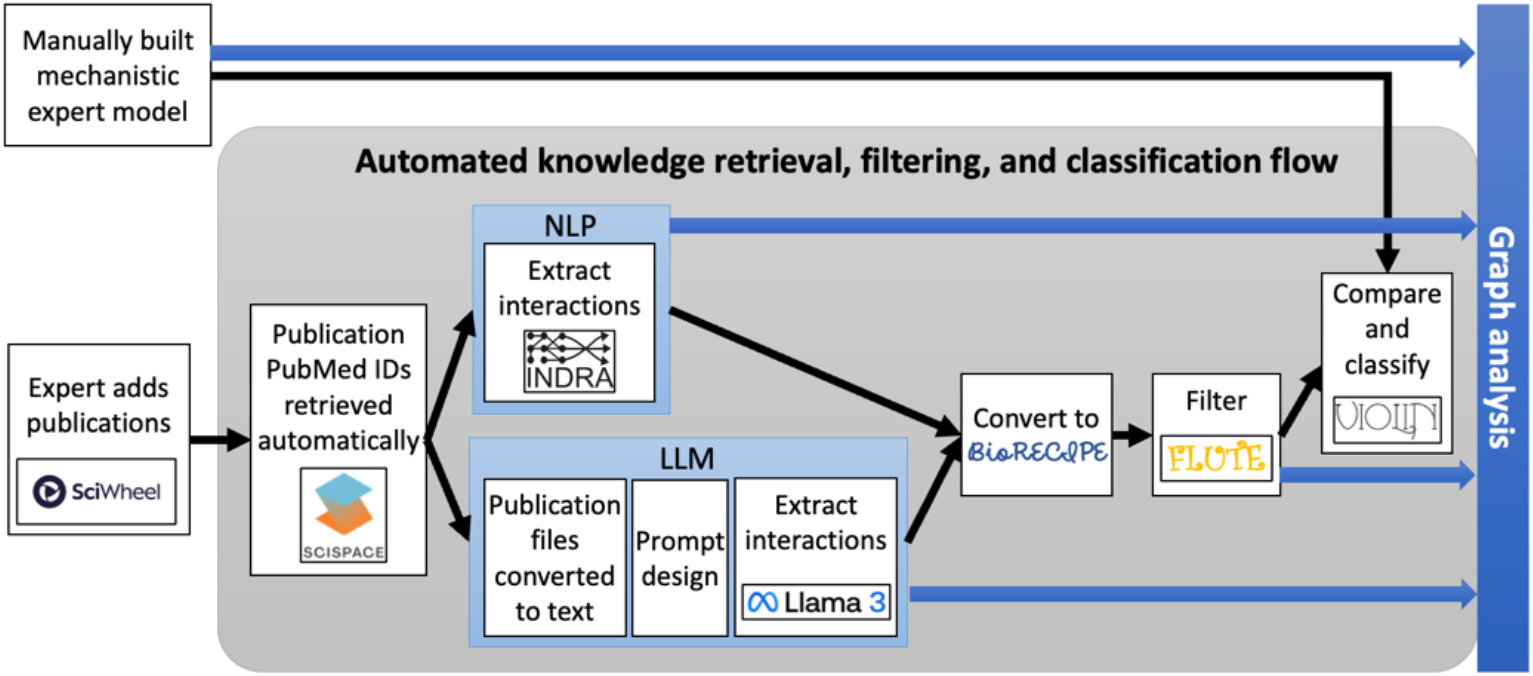
Our automated pipeline for collecting new knowledge from literature and evaluating NLP and LLM approaches with interaction filtering and classification tools, and with graph metrics.

For instance, while INDRA [6] excels at accessing a broad range of sources to identify interactions, it has limitations in retrieving certain papers. On the other hand, LLMs can access and process these missing papers to uncover additional interactions. By combining these approaches, we were able to identify more interactions and address the gaps in each method. The first approach utilizes the INDRA framework, a computational tool that extracts relevant information from research papers and organizes it in a structured JSON format. INDRA integrates NLP methods and tools such as REACH [7] and TRIPS [8], and interaction databases, such as Signor [9], BioGRID [10], and BioPAX [11]. For the LLM-based approach, we used LLaMa 3 [12] and created several scripts to convert papers to plain text format, access LLaMa through an API, and instruct it how to output the collected information with few-shot prompting. The information extracted from literature by INDRA, LLaMa is then converted into a structured list of interactions in a tabular BioRECIPE format [13].

The interaction lists are also filtered with FLUTE [14] to keep only those that are highly supported by interaction databases. Although this step increases the confidence in the interaction list at the output of FLUTE, it may remove more recent and novel observations that are not yet included in databases, and which have a potential to improve the model. Therefore, we conducted our analysis of the interactions lists both before and after the FLUTE filtering step.

We compared these interaction lists with the expert-built model using VIOLIN [15], which classifies them into four categories, and within each category into several sub-categories. VIOLIN finds in the interaction lists those interactions that corroborate or contradict the model, the interactions that can extend the model, and the interactions that require further investigation (potentially erroneous outputs from NLP/LLM). Biological networks exhibit certain properties by nature, such as being scale-free. Investigating these network properties helps identify key features. So, we used several graph metrics available in Cytoscape [16] to investigate the differences between the structure of the manually built model and the knowledge automatically extracted with the traditional NLP or new LLM approaches. Similarly, analyzing properties in our GBM model can provide valuable insights and guide discussions with experts.

## III. Results and Discussion

The Sankey diagram in Figure 2(a) shows the flow of collecting and processing papers in several stages. In Figure 2(b), we present summary statistics of several networks, GBM model, INDRA output before and after filtering with FLUTE, and LLaMA output before and after filtering with FLUTE. The GBM model has by far the largest network diameter (30) indicating that it allows for signal transduction on longer pathways. INDRA extracted 5,297 interactions; however, the high number of connected components (1,154) indicates a highly fragmented network with many isolated groups. Even though it has a large number of interactions, the diameter of the network formed by INDRA’s output is only half of the GBM model diameter. FLUTE significantly refined the network from INDRA, reducing the number of connected components by approximately 92%, resulting in a more unified structure. LLaMa generated a smaller set of 207 interactions, yet its output network was also very disconnected, with 160 separated clusters. The GBM model also has several clusters following expert recommendation to add specific elements despite limited knowledge on their connections to the rest of the network. All networks display low clustering coefficients, indicating weak local interconnectedness. After filtering, the network density increased, with 420 and 23 interactions remaining in INDRA and LLaMA output, respectively. We also explored the distribution of element in these networks across several element types: genes, RNAs, proteins, protein families, chemicals, biological processes, and other (Figure 2(c)). The gene, RNA, and protein categories (protein family is merged within protein category) are equally distributed in the GBM model, and there are very few chemicals and several biological processes. INDRA, on the other hand seems to output mainly proteins and chemicals, while more than 80% of elements are considered as “other” and likely many are machine reader errors. LLaMA also finds mainly proteins and a larger percent of genes, chemicals and biological processes than INDRA, while almost half the elements it finds are under “other” category.

**Figure 2.**
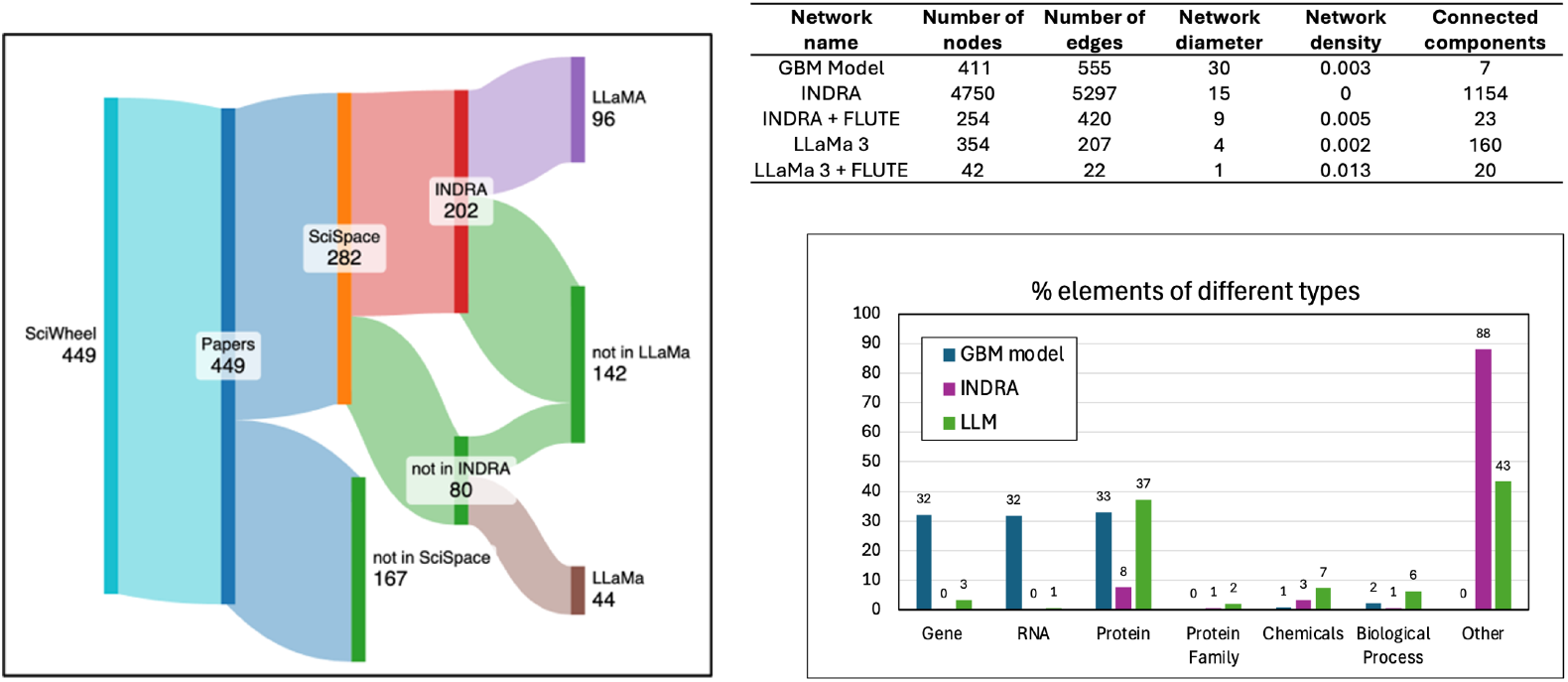
(a) Sanky diagram indicating papers processed and the number of interactions obtained. From the expert’s collection of 449 papers, we were able to identify only 282 paper IDs using SciWheel. The paper IDs were then input into INDRA, which successfully identified 202 papers. To use LLaMa, we collected papers from PubMed through an API, which allowed access only to 140 out of 282 papers. Of these 140 papers, LLaMa was able to process 44 out of 80 papers that INDRA could not find (b) Graph characteristics for the GBM model, the networks obtained using INDRA (traditional NLP-based flow) or LLaMa 3, and for the networks after the INDRA and LLaMa 3 outputs filtered with FLUTE. (c) The distribution of different element types in the GBM model, INDRA output, and LLaMa 3 output.

Figure 3 shows the output from VIOLIN, classification of interactions obtained from INDRA and LLaMA with respect to the GBM model. The blue pie charts indicate the total number of corroborations found in these two interaction lists, distributed across several corroboration sub-categories. None of the interactions in these two lists was a strong match to the model. INDRA found seven indirect interactions that matched model interactions and 20 interactions that matched paths in the model, while LLaMa found only one interaction that added new information to an existing model interaction. The green pie charts show the number of interactions that can be used to extend the model. Most of the interactions from the INDRA and LLaMa output belong to this category. In the INDRA output, there are 3,677 full extensions representing entirely new, disconnected interactions; 1,189 hanging extensions indicating new interactions where only one node already exists in the model; and 203 internal extensions suggesting potential refinements within the network, as both nodes exist in the model but are currently unconnected. Extensions in LLaMa’s output are similarly distributed across sub-categories, with most being full extensions, a substantial fraction of hanging extensions and several internal extensions. Interestingly, only one contradiction is found in INDRA’s output, while five contradictions are in LLaMA’s output (yellow charts). That could potentially indicate LLaMA’s issue with hallucination. On the other hand, VIOLIN did not flag many of LLaMa’s interactions for further investigation, while more interactions were flagged in INDRA’s output (red charts).

**Figure 3.**
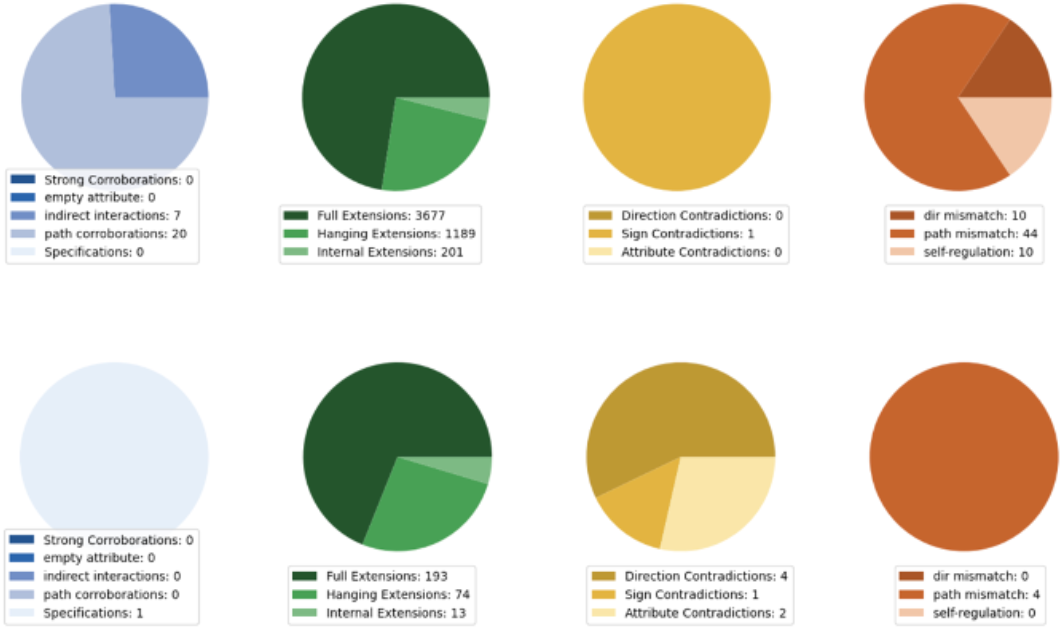
Classification of interactions in outputs from INDRA (top) and LLaMA (bottom) with respect to the GBM model. Interactions are classified into four categories: corroborations (blue), extensions (green), contradictions (yellow) and flagged (red). Each category also has several sub-categories.

Using Cytoscape we also investigated the distribution of individual nodes in the GBM model, INDRA and LLaMa outputs with respect to several graph metrics (Figure 4): average shortest path length, betweenness centrality, closeness centrality, clustering coefficient, in-degree, out-degree, stress, and eccentricity. These metrics confirm our expectation that, even though the selection of literature was very focused and expert-guided, the interaction networks obtained from papers have highly disjoint nature. Our next steps will include identifying paths of connected interactions in NLP/LLM output, an exploration of other literature selection strategies, and an in-depth investigation of contradictions, extensions, and flagged interactions that VIOLIN identified with knowledge of experts.

**Figure 4.**
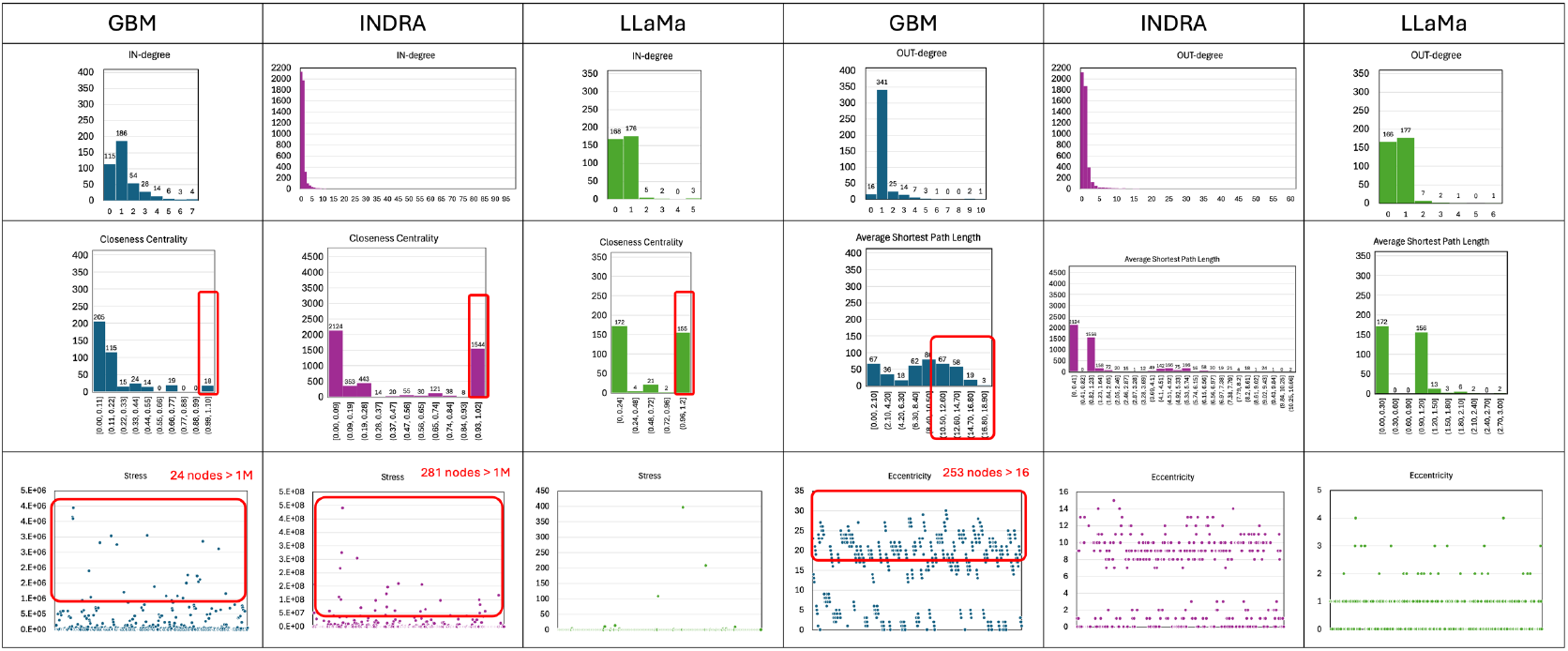
Several graph features meatured with the GBM model, INDRA output, and LLaMa output. Top two rows show distributions of metric values for each network. The last row shows the metric value for each node in the network.

## Acknowledgements

This projects was funded in part by the NSF EAGER Award #2324742.

## Notes

### Competing Interest Statement

The authors have declared no competing interest.

## References

[1] E. Holtzapple, B. Cochran, and N. Miskov-Zivanov, “Automated verification, assembly, and extension of GBM stem cell network model with knowledge from literature and data,” bioRxiv, 2021.

[2] E. Holzapple, N. Miskov-Zivanov, and B. Cochran, “CSIG-13. A DYNAMIC CAUSAL MODEL OF GLIOBLASTOMA STEM CELL SIGNALING PREDICTS EFFECTS OF KINASE INHIBITORS,” Neuro Oncology, vol. 23, 2021.

[3] Y. Ahmed, C. A. Telmer, and N. Miskov-Zivanov, “CLARINET: efficient learning of dynamic network models from literature,” Bioinform Adv, vol. 1, no. 1, p. vbab006, 2021.

[4] Y Ahmed, CA Telmer, G Zhou, and N. Miskov-Zivanov, “Context-aware knowledge selection and reliable model recommendation with ACCORDION,” Frontiers in Systems Biology, vol. 4, 2024.

[5] Sciwheel. “Sciwheel – A Reference Management and Research Collaboration Tool.” (accessed 2025).

[6] B. M. Gyori, J. A. Bachman, K. Subramanian, J. L. Muhlich, L. Galescu, and P. K. Sorger, “From word models to executable models of signaling networks using automated assembly,” Molecular Systems Biology, vol. 13, no. 11, p. 954, 2017.

[7] M. A. Valenzuela-Escárcega, G. Hahn-Powell, and M. H. Surdeanu, T., “A Domain-independent Rule-based Framework for Event Extraction,” presented at the ACL-IJCNLP 2015 System Demonstrations, Beijing, China, 2015.

[8] G. A. Ferguson, James F., “TRIPS: An Integrated Intelligent Problem-Solving Assistant,” presented at the Proceedings of the Fifteenth National Conference on Artificial Intelligence (AAAI-98), Madison, Wisconsin, USA, 1998.

[9] P. Lo Surdo et al., “SIGNOR 3.0, the SIGnaling Network Open Resource 3.0: 2022 Update,” Nucleic Acids Research, vol. 51, no. D1, pp. D631–D637, 2023.

[10] R. Oughtred et al., “The BioGRID database: A comprehensive biomedical resource of curated protein, genetic, and chemical interactions,” Protein Science, vol. 30, no. 1, pp. 187–200, 2021.

[11] E. Demir et al., “The BioPAX community standard for pathway data sharing,” Nat Biotechnol, vol. 28, no. 9, pp. 935–42, Sep 2010.

[12] L. T. A. a. Meta, “The Llama 3 Herd of Models,” arXiv preprint, 2024.

[13] E. Holtzapple et al., “The BioRECIPE Knowledge Representation Format,” ACS Synth Biol, vol. 13, no. 8, pp. 2621–2624, Aug 16 2024.

[14] E. Holtzapple, C. A. Telmer, and N. Miskov-Zivanov, “FLUTE: Fast and reliable knowledge retrieval from biomedical literature,” Database (Oxford), vol. 2020, Jan 1 2020.

[15] H. Luo et al., “Context-driven interaction retrieval and classification for modeling, curation, and reuse,” bioRxiv, 2024.

[16] P. Shannon et al., “Cytoscape: a software environment for integrated models of biomolecular interaction networks,” Genome Res, vol. 13, no. 11, pp. 2498–504, Nov 2003, doi: 10.1101/gr.1239303.

